# Interoceptive Awareness of the Breath Preserves Dorsal Attention Network Activity amidst Widespread Cortical Deactivation: A Within-Participant Neuroimaging Study

**DOI:** 10.1101/2022.05.27.493743

**Authors:** Norman A. S. Farb, Zoey Zuo, Cynthia J. Price

## Abstract

Interoception, the representation of the body’s internal state, serves as a foundation for emotion, motivation, and wellbeing. Yet despite its centrality in human experience, the neural mechanisms of interoception are poorly understood. The Interoceptive/Exteroceptive Attention Task (IEAT) is a novel neuroimaging paradigm that compares behavioral tracking of the respiratory cycle (Active Interoception) to tracking of a visual stimulus (Active Exteroception). Twenty-two healthy participants completed the IEAT during two separate scanning sessions (N = 44) as part of a randomized control trial of Mindful Awareness in Body-oriented Therapy (MABT). Compared to Exteroception, Interoception deactivated somatomotor and prefrontal regions. Greater interoceptive sensibility (MAIA scale) predicted sparing from deactivation along the anterior cingulate cortex (ACC) and left-lateralized language regions. The right insula—typically described as a primary interoceptive cortex—was only implicated by its further deactivation during an exogenously paced respiration condition (Active Matching). Psychophysiological interaction analysis characterized Active Interoception as promoting greater ACC connectivity with lateral frontal and parietal regions commonly referred to as the Dorsal Attention Network. Interoception of the breath may therefore involve reduced cortical *activity* but greater *connectivity*, with greater sensibility sparing cortical inhibition within well-characterized attentional networks. In contrast to a literature that relates detection of liminal signals such as the heartbeat to anterior insula activity, attention towards accessible body sensations such as the breath may lead to a low activity, high connectivity state in which sensory signals from the body may be better discerned.

**Significance Statement:** Interoception, the representation of the body’s internal state, is poorly understood compared to the external senses, with existing neuroimaging studies failing to match task difficulty between interoceptive and exteroceptive tasks. The present study used a novel fMRI task to compare interoceptive and exteroceptive attention, and how this distinction was moderated by self-reported interoceptive awareness. The results implicate three novel interoceptive mechanisms: interoception reduces cortical *activity* while increasing *connectivity*, wherein awareness is linked to preserved activation of the brain’s salience network and left-lateralized language regions. These findings characterize interoception as a lower activity state in which awareness depends upon the ability to notice and report on body signals typically obscured by the processing of exteroceptive information and other forms of cognition.

## Introduction

Interoception, the sense of the body’s internal state, is central to human experience, providing homeostatic cues (Strigo & Craig, 2016) that inform emotion (Barrett, 2017; Wiens, 2005), motivation (Craig, 2003; Critchley & Garfinkel, 2017), and wellbeing (Chen et al., 2021; Farb et al., 2015; Paulus & Khalsa, 2021). Yet interoception’s neural mechanisms are still poorly understood when compared to the five “canonical” human senses (Chen et al., 2021).

Our understanding has been limited by a focus on interoceptive *accuracy* by the research community. For example, heartbeat detection paradigms model the ability to detect a liminal cardiac signal (e.g., Garfinkel et al., 2015). Superior detection accuracy is linked to greater activation of the anterior cingulate (ACC) (Critchley et al., 2004) and anterior insula (Chong et al., 2015), hubs of the brain’s “Salience Network” (SLN) (Chand & Dhamala, 2016; Seeley et al., 2007). Yet SLN recruitment may indicate a broader error-monitoring system that does not distinguish interoception from other sensory processes (Baltazar et al., 2021).

Moreover, differences in interoceptive accuracy may not be especially relevant for wellbeing. Clinical populations often show normal interoceptive accuracy (Desmedt et al., 2020), and clinically-efficacious interoception-focused interventions such as mindfulness training do not improve accuracy (Khalsa et al., 2020; Parkin et al., 2014). Patients with anxiety or panic disorders have historically demonstrated *superior* accuracy (Ehlers & Breuer, 1992; Zoellner & Craske, 1999), but tend to catastrophize interoceptive experience (Domschke et al., 2010). The complex relationship between interoceptive accuracy and wellbeing demonstrates the need to expand the scope of interoceptive research to other, more clinically relevant constructs.

The ability to sustain interoceptive *attention* remains a compelling if underexplored determinant of mental health. Improving the capacity to skillfully attend to interoceptive cues remains a central target of contemplative interventions such as mindfulness training (Gibson, 2019; Price & Weng, 2021). Such interventions address intolerance for interoceptive signals, which otherwise leads vulnerable individuals into patterns of experiential avoidance (Anestis et al., 2007; Hayes et al., 1996). Interoceptive *sensibility —* how one appraises interoceptive signals — shows stronger links to wellbeing than interoceptive accuracy (Ferentzi et al., 2019; Schuette et al., 2021), with a growing literature relating trust in interoceptive signals to greater wellbeing (Calì et al., 2015; Trevisan et al., 2019).

Interoceptive attention seems to recruit a distinct neural network from exteroception that features the middle insula (Haruki & Ogawa, 2021), consistent with its proposed role as a bridge to the prefrontal cortex from primary representation cortices in the posterior insula and somatosensory regions (Craig, 2009; Farb et al., 2013a; Pollatos et al., 2007). Activity in the middle insula characterizes wellbeing (Nord et al., 2021), with hypoactivation linked to depression (Avery et al., 2014; Farb et al., 2010; Farb et al., 2022), and hyperactivation linked to anxiety (Kerr et al., 2016; Tan et al., 2018). Breath monitoring in particular appears to strengthen coherence across a frontotemporal-insular network (Herrero et al., 2018).

Yet neuroimaging studies characterizing interoceptive attention to the breath have been confounded by non-equivalent task demands. Farb et al. (2013) contrasted exteroceptive and interoceptive attention by comparing a visual condition requiring behavioral responses against passive breath monitoring; Wang et al. (2019) compared a more difficult interoception task against an easier exteroception task. In both cases, the more demanding task recruited the Dorsal Attention Network (DAN), while the easier/passive task implicated the Default Mode Network (DMN), confounding conclusions on interoceptive representation.

We therefore developed a novel fMRI paradigm for characterizing the neural dynamics of breath-focused interoception vs. a more closely balanced exteroception task. We hypothesized (H1) that interoception would result in greater somato-insular recruitment than exteroception, but reduced DAN activation. We further hypothesized (H2) that greater self-reported interoceptive sensibility would correlate with greater SLN activity during interoception, consistent with a greater capacity for interoceptive monitoring and integration. We also explored: (H3) a paced-breathing condition to understand impact of endogenous vs. exogenous respiratory control, and (H4) functional connectivity differences between Interoception and Exteroception.

## Materials and Methods

### Experimental Design

This fully within-participant study was conducted to validate a novel interoceptive attention task as part of an NIH-funded pilot study, a two-group randomized control trial to examine the neural correlates of interoceptive awareness in the context of Mindful Awareness in Body-oriented Therapy (MABT) training (www.ClinicalTrials.gov identifier: NCT03583060). MABT is a well-validated, clinical intervention the focuses on developing the interoceptive capacities of identifying, accessing, and appraising internal bodily signal to support adaptive emotion regulation (Price & Hooven, 2018). To maximize power, data from the study’s two assessment timepoints were combined into a single dataset, with group and time included in all statistical models as nuisance covariates.

### Participants

Twenty-two right-handed study participants (11 male and 11 female), of adult age (mean: 36.1 years, range: 18-62) completed both baseline and post-intervention assessments. Eleven participants (50% of the sample) were randomly allocated to receive eight MABT sessions, delivered individually once per week for eight weeks. Twenty participants self-identified as Caucasian, one as African American, and two as Hispanic. Their highest education levels were high school (n = 5), 2 years of college (n = 2), Bachelor’s degree (n = 8), and Master’s degree or higher (n = 7).

Sample size for the study was determined by simulation-based power analysis. Given the difficulties in *a priori* registration of all fMRI contrasts, power analysis was conducted to determine minimum sample size for up to 10 focal contrasts while maintaining familywise power ≥ .90; this required a per-test power of 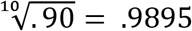. The alpha level was determined by taking the typical peak Z required for peak voxel familywise error correction (Z = 4.53), computing its equivalent *p*-value from the normal distribution (*p* = 2.95 x 10^-6^), and then Bonferroni correcting this value for 10 comparisons (*p* = 2.95 x 10^-7^). The simulated effect size was determined by computing the average of the peak Z scores from all significant clusters revealed by the [Interoception – Exteroception] in Farb et al. (2013), as were the variances of the contrast main effect and the participant random effects (Z = 5.87, *s^2^_contrasteffect_* = .20, *s^2^_randomeffect_* = .22). Monte Carlo simulation conducted in the R statistical environment suggested that up to 10 contrasts would be sufficiently powered at N ≥ 9 (Figure S1), so the parent trial sample size of N = 44 scans adequately powered the study design.

Twenty-five healthy individuals with self-reported elevated stress were initially recruited through advertisements in a local newspaper and through the University of Washington (UW) research volunteer website and flyers posted on the UW campus. Inclusion criteria were: 1) being over 18 years of age; 2) Perceived Stress Scale (Taylor, 2015) scores indicating moderate stress levels; 3) naive to mindfulness-based approaches (no prior experience), 4) agrees to forgo (non-study) manual therapies (e.g., massage) and mind-body therapies (e.g., mindfulness meditation) for 12 weeks (baseline to post-test); 5) fluent in English; 6) can attend MABT and assessment sessions; and 7) right-handed (for uniformity of neuroimaging results). Exclusion criteria were: 1) lifetime diagnosis of mental health disorder; 2) unable to complete study participation (including planned relocation, pending inpatient treatment, planned extensive surgical procedures, etc.); 3) cognitive impairment, assessed by the Mini-Mental Status Exam (MMSE) (Folstein et al., 1975) if demonstrated difficulty comprehending the consent; 4) use of medications in the past 30 days that affect hemodynamic response; 5) lifetime head injuries or loss of consciousness longer than 5 minutes; 6) currently pregnant; or 7) contraindications for MRI, e.g., claustrophobia, metal objects in body, etc. The full CONSORT diagram is available in the supplementary materials (Figure S2).

### Ethics Statement

All participants provided informed consent. The study procedures were reviewed and approved by the institutional review board at the University of Washington in accord with the World Medical Association Declaration of Helsinki.

### The Interoceptive/Exteroceptive Attention Task (IEAT)

The Interoceptive/Exteroceptive Attention Task (IEAT) is a novel paradigm for exploring the neural dynamics of respiratory attention and awareness. The IEAT consisted of five conditions: Passive Exteroception, Passive Interoception, Active Interoception, Active Exteroception, and Active Matching (a paced breathing condition), as shown in Figure 1.

**Figure 1.**
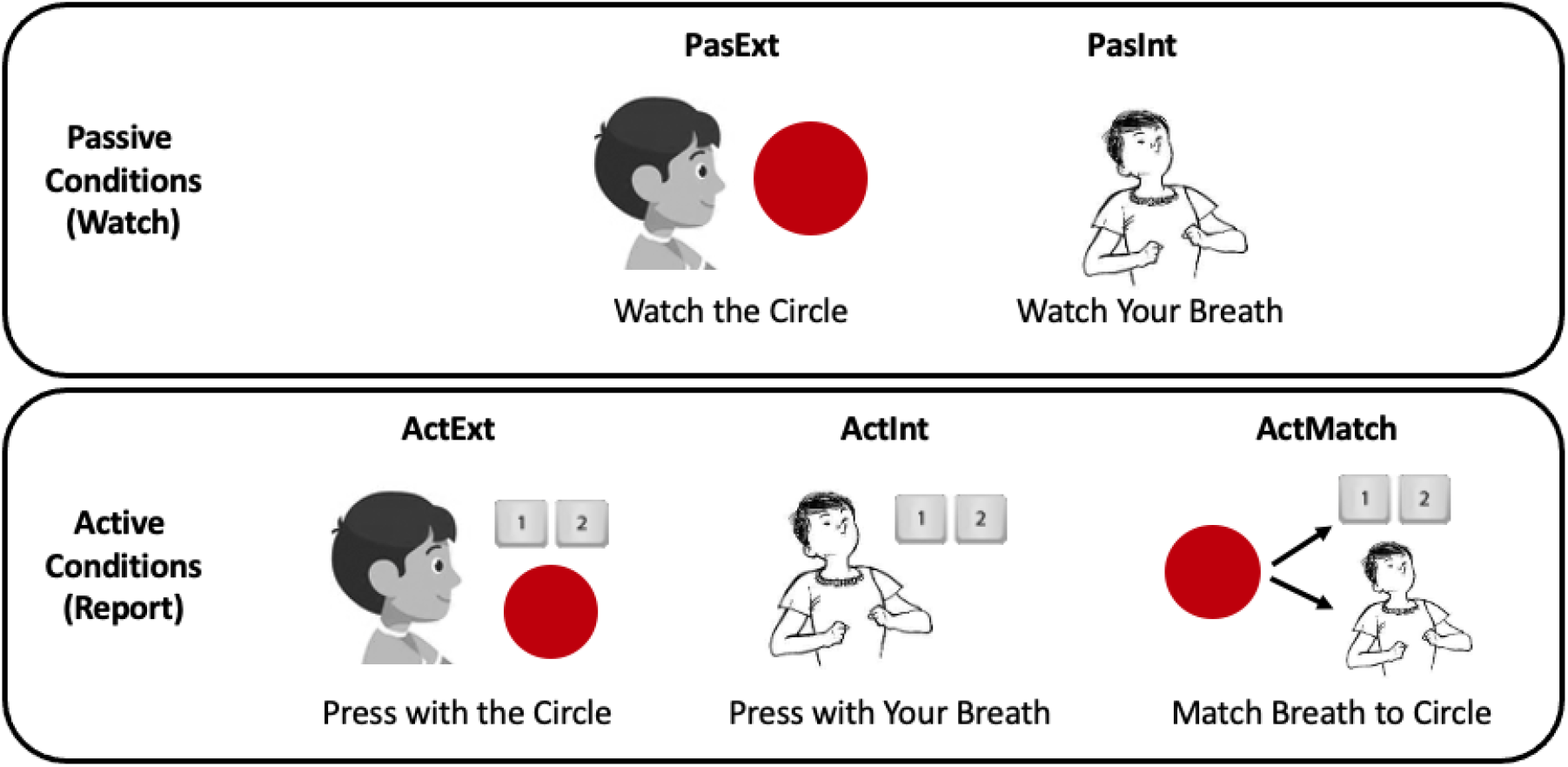
Schematics for the Interoceptive/Exteroceptive Attention Task (IEAT). In Exteroceptive conditions, participants attended to a circle expanding and contracting; in Interoceptive conditions, participants attended to their respiratory inhalation and exhalation. In Passive conditions, participants simply observed the circle or their breath; in Active conditions, participants pressed buttons to track the circle or breath. In the Matching condition, participants tracked the circle’s movements while synchronizing their respiration cycle with the circle.

#### Passive Conditions

During Passive Exteroception, participants were asked to visually monitor a circle as it expanded and contracted periodically on the MRI-compatible visual display without making any behavioral responses. The circle’s pulse frequency was set to match participants’ in-scanner breathing frequency, obtained from a respiration belt worn by participants in the scanner (usually around 12 Hz). During Passive Interoception, participants viewed a stationary circle on the screen while attending to sensations of the breath.

#### Active Conditions

During Active Exteroception, participants reported on the expansion and contraction of the circle on the screen, which again was set to pulse at participants’ in-scanner respiratory frequency. During Active Interoception, participants reported on inhalations and exhalations by making button box key presses with their right-hand index and middle fingers respectively. The circle on the screen expanded and contracted with these key presses, approximating the frequency of circle movement during Passive Exteroception. During Active Matching, participants reported on the expansion and contraction of the circle as in Active Exteroception, while synchronizing their inhalation to the circle’s expansion and their exhalation to the circle’s contraction.

### Self-Reported Interoceptive Awareness

Perhaps the most common and general (i.e., not task-specific) index of subjective interoceptive engagement is the Multidimensional Assessment of Interoceptive Awareness (MAIA), which in validation has demonstrated strong associations to subjective wellbeing (Mehling et al., 2012, 2018). The MAIA is a broad and general measure that examines multiple indicators of subjective interoceptive attention including awareness, comfort, skill at sustaining attention, and trust. The MAIA is sensitive to treatment effects from clinically efficacious interoceptive-focused interventions, with studies demonstrating a positive relationship between improved interoceptive awareness on the MAIA and treatment health outcomes (Fissler et al., 2016; Price et al., 2019; Roberts et al., 2022).

Here, the MAIA was used to provide a subjective report of the ability to adaptively engage interoceptive attention. The MAIA is a 32-item scale designed to measure multidimensional facets of self-reported interoceptive awareness relevant for wellbeing. The eight subscales canvas experiential domains such as Noticing, Listening, and Emotional Awareness, as well as regulatory domains such as Attention Regulation, Self-Regulation, and Trusting. As noted above, the MAIA has an updated version to improve the reliability of the two reverse-coded subscales (Mehling et al., 2018), but as the new version was not available at study outset, the total score from the original version was employed, which yielded very good reliability (Cronbach’s α = .85) in the present sample.

### Procedures

Participants completed the IEAT during fMRI acquisition at two timepoints, baseline and 3-month follow-up. The MAIA self-report questionnaire was administered to assess interoceptive awareness at both timepoints. Training effects are the subject of separate reports.

### Data Analysis

#### Respiration Confounds

Changes in breathing depth and rate modulate CO2 concentration in the bloodstream, with faster, shallower breathing increasing blood oxygen level dependent (BOLD) signals throughout the brain (Birn et al., 2006). Failure to correct for task-related variation in breathing rate and depth can therefore lead to spurious associations with experimental conditions (Weiss et al., 2020). Consensus on correction for such effects is an area of ongoing investigation, but fMRI analysis should at a minimum correct for respiratory frequency and volume / time (Power et al., 2020).

#### Respiration Frequency

Respiration data was acquired using a MR-compatible respiration belt sampling at 500 Hz. Respiration data was first smoothed using a 1-sec zero phase low-pass filter window and then mean-corrected. Breath frequency was then estimated using a Fast Fourier Transform (FFT) of the respiration period. As a first level covariate, frequency was estimated using a 10-sec sliding window across the time series, generating a frequency value for each volume acquired in the timeseries. Trial-specific frequencies were also estimated across each task period to serve as covariates at the second (group) level of analysis.

#### Respiration Volume / Time

The respiratory signal is influenced by changes in respiratory volume in addition to frequency, with Respiratory Volume per Time (RVT) predicting widespread changes in BOLD activity (Birn et al., 2006). To correct for this influence, RVT was calculated as recently recommend (Power et al., 2020) by first finding peaks and troughs of the normalized (z-scored) respiratory signal, setting a minimum distance between peak and trough of 0.5 standard deviations, but a minimum distance of 2 seconds between consecutive peaks or troughs. Linear interpolation of peaks and troughs generated an ‘envelope’ around the respiratory signal, and the difference between peak and trough values represented the RVT score across the timeseries. Trial-averaged RVTs were also estimated for each task period for use as covariates at the second (group) level of analysis.

#### Stimulus and Behavior Timeseries

The three active localizer conditions, Active Exteroception, Active Interoception, and Active Matching required button-presses to track the sensory target, i.e., inhalation/exhalation during the respiratory cycle, or expansion/contraction during the visual circle cycle. Each active condition therefore produced three periodic timeseries: (i) the circle radius, as measured by timestamps in the participant log files, (ii) respiration volume, as measured by pressure transduction on an MR-compatible respiration belt, and (iii) a waveform generated from participant button presses, with button presses indicating inflection points (peaks and troughs) of the response waveform.

The respiration belt waveform was obtained directly from a physiological logfile for each fMRI run. Respiration data was sampled at 500 Hz and included a scanner-generated timestamp to precisely indicate the beginning and end of the functional run. Respiration data was segmented into task trials using timestamps from participant logfiles. The ‘pracma’ library (Borchers, 2021) in the R statistical programming environment (R Core Team, 2017) was used to find peaks and troughs within the timeseries (Figure 2A below). The minimum peak difference was set to the sampling rate, as individual breaths were unlikely to have a period shorter than 1 second.

**Figure 2.**
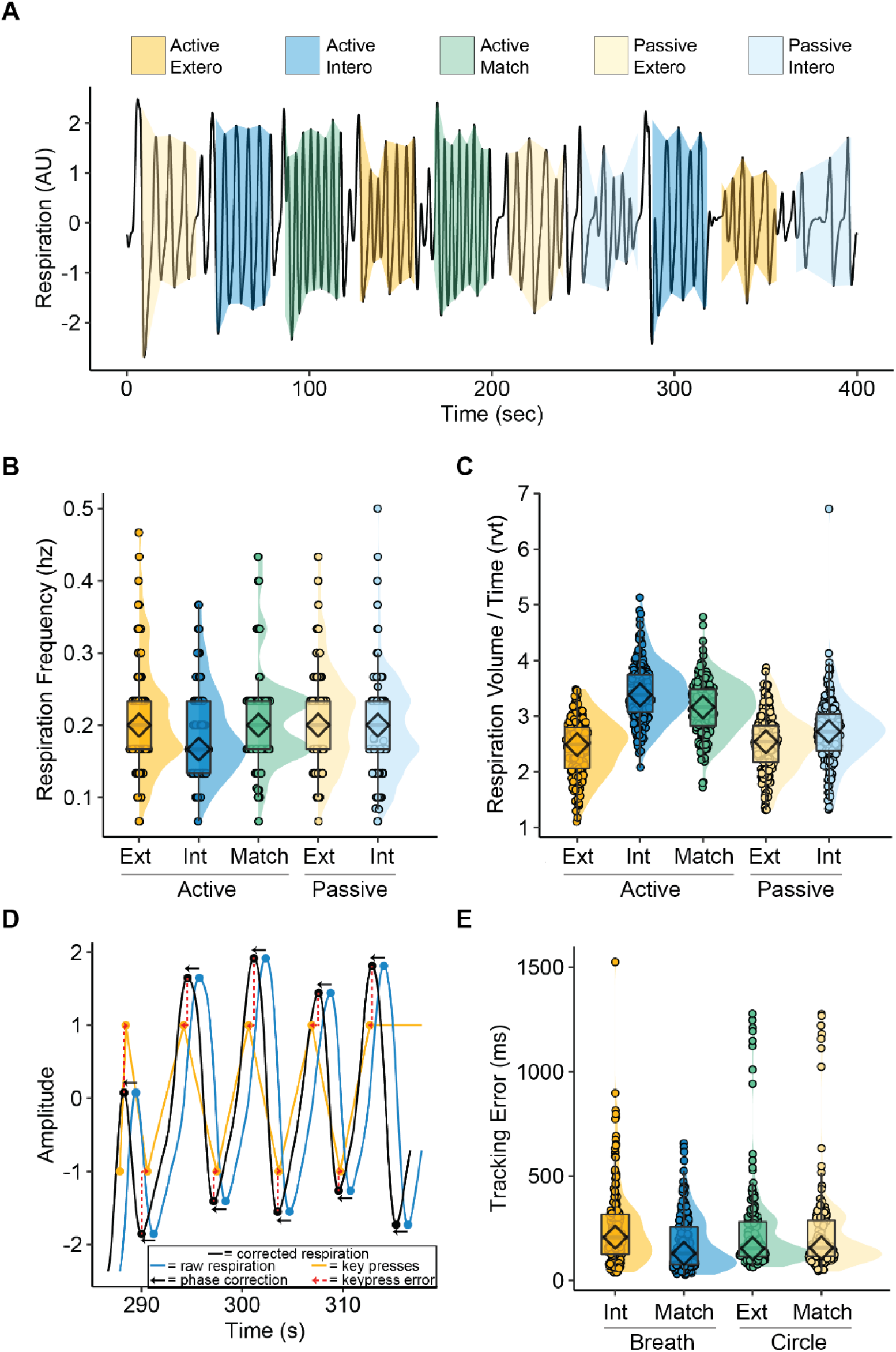
Respiration and Behavior Tracking. **(A)** Respiratory peak and trough detection was successful for each of the experimental task conditions, allowing for calculation of respiratory frequency (Hz) and respiratory volume over time (RVT). **(B)** Respiration was slower during Active Interoception than in other conditions. **(C)** Respiration was deeper (greater volume / time) during Active Interoception and Active Matching than in other conditions. **(D)** To assess tracking accuracy, the respiration waveform (blue) was first temporally adjusted (black) to match phase with keypress signals (orange). The average error (red) between key press timing and respiratory peaks/troughs was calculated; the same procedure was applied to a waveform describing circle radius in the exteroceptive tracking conditions. **(E)** No differences in tracking accuracy were observed between Active Interoception of the Breath and Active Exteroception of the Circle, nor with but Active Matching to the Circle; however, breath tracking was superior during the Active Matching condition.

Behavioral keypress waveforms were generated using linear interpolation of behavioral logfile timestamps for button presses using the *approx* function in R. The inhalation/expansion button “1” was coded as −1 to indicate beginning from the trough of the period, and the exhalation/contraction button “2” was coded as 1 to indicate peak of the period. As the waveforms were intended to be compared to the respiratory signal, the waveforms were interpolated at a sampling rate of 500 Hz to match the data acquired by the respiration belt.

The circle stimulus waveform was obtained from the stimulus presentation software. The circle period was matched to participant-specific breathing frequencies obtained via respiration belt during structural MRI acquisition. For each trial, the timeseries began at an arbitrary unit of 0. The expansion/contraction period was symmetrical and defined as half the participant respiratory period. The circle value was then extrapolated as increasing from −1 for the expansion period to reach a value of 1 by the end of the period; the circle then paused for 0.2 * the expansion period; the circle then decreased at the same rate for the contraction period to reach a value of −1 by the end of the period, and then paused again for 0.2 * the contraction period, repeating the cycle the end of the task block.

#### Tracking Accuracy

To calculate trial-specific tracking accuracy in the active tracking conditions (Active Interoception, Active Exteroception, and Active Matching), the appropriate sensory stimulus waveform (respiration in Active Interoception, the visual circle in Active Exteroception) was first phase-corrected to maximally align with the keypress waveform; this correction served to compensate for systematic signal transduction lags between the three types of signals. Phase correction was performed using a normalized cross-correlation (NCC) analysis using the ‘dtwclust’ library in R to estimate the maximum correlation between behavior and respiration/circle timeseries (Sardá-Espinosa, 2019). The NCC analysis tested correlation as function of lag between two timeseries, up to a full second (500 samples) of lag.

Following phase correction, tracking accuracy was assessed by computing the deviation from the respiration/circle inflection points to the nearest keypress, as tracking keypresses were cued by changes in circle/respiration phase (the switch from expansion/inhalation to contraction/exhalation). The average deviation in ms was then calculated for each trial and used as a measure of tracking accuracy, with lower scores indicating more accurate tracking (Figure 2D below). The complete algorithm for generating timeseries waveforms and calculating tracking accuracy is available on the Open Science Framework (https://osf.io/cuz6s).

#### Neuroimaging Data Acquisition

Neuroimaging was performed using a 3T Philips Achieva scanner (Philips Inc., Amsterdam, Netherlands) at the Diagnostic Imaging Sciences Center, University of Washington. Imaging began with the acquisition of a T1-weighted anatomical scan (MPRAGE) to guide normalization of functional images (~ 6 minutes) with TR= 7.60 ms, TE = 3.52 ms, TI = 1100 ms, acquisition matrix = 256 × 256, flip angle = 7°, shot interval = 2530 ms, and 1mm isotropic voxel size. Functional data were acquired using a T2*-weighted echo-planar-imaging (EPI) sequence with TR = 2000, TE = 25 ms, flip angle α = 79°, field of view = 240 × 240 × 129 mm, 33 slices, and a voxel size of 3 × 3 × 3.3 mm with 3.3 mm gap. Button presses were registered using a 2-button MR-compatible response pad.

#### Preprocessing

A set of preprocessing steps was performed using the consortium-developed fMRIprep robust preprocessing pipeline for fMRI data (https://fmriprep.readthedocs.io/en/stable/). Briefly, preprocessing consisted of realignment and unwarping of functional images, slice timing correction, and motion correction. The functional images were resliced using a voxel size of 2 × 2× 2 mm and smoothed using a 6-mm FWHM isotropic Gaussian kernel.

#### First Level Analysis

Within subject statistical models were used to characterize the interoceptive networks at the participant level. Participant time series from the interoceptive tasks were submitted to separate first-level general linear statistical models using Statistical Parametric Mapping software (v12). Task-specific boxcar stimulus functions were convolved with the canonical hemodynamic response function, separately modeling the onsets of the interoceptive and visual control conditions for each participant. To control for motion and physiological confounds, six standard movement parameters, the root mean square of temporal change (DVARS), framewise displacement, respiration rate, and RVT were all included as nuisance covariates.

#### Second Level Analysis

Participant first-level maps for each experimental condition [Passive Interoception, Passive Exteroception, Active Interoception, Active Exteroception, Active Matching] were analyzed at the second level using a full-factorial mixed-model ANOVA in SPM12 (Friston, 2007). The second level contrasts first subdivided the tasks in two main effects and their interaction term: Reporting Demand [Active vs. Passive] × Target [Interoception vs. Exteroception]. Follow up comparisons within Active [Active Interoception vs. Active Exteroception vs. Active Matching] and within Passive [Passive Interoception vs. Passive Exteroception] were also modelled.

Familywise control for multiple comparisons (corrected *p* < 0.05) in whole-brain analyses was achieved through threshold-free cluster enhancement (TFCE), which controls familywise error rate based on a permutation testing approach and determines optimal voxel-wise cluster-forming thresholds using an automated algorithmic method (Smith & Nichols, 2009). The algorithm eliminates the need to choose between arbitrary correction thresholds by sampling across statistically equivalent peak and cluster size thresholds. Trial-averaged respiration frequency and RVT were modelled at the second level as nuisance covariates.

#### Trial-level Confounds

The current study comes from an exploratory clinical trial, the results of which are the subject of a separate report. We combined data across the trial to power the comparison of IEAT task conditions and modelled any effects of trial Group (MABT vs. Control), Time (Baseline vs. Post-Intervention) and their interactions as nuisance covariates. All models also contained condition-averaged respiration rate as a covariate, to further control for variation in respiration between experimental conditions. Post hoc analyses that did not include the nuisance covariates of Group and Time did not meaningfully change the reported results.

#### Region of Interest (ROI) Analysis

For region of interest (ROI) analysis, all signal extractions were taken from models containing the nuisance covariates. Using the built-in SPM12 function, the median value of the raw, unwhitened signal was extracted from all voxels within the ROI, yielding one value per participant at each time point. These values were entered into a linear mixed-effects model with restricted likelihood estimation was applied using the ‘lme4’ library (Bates et al., 2015) in the R statistical programming environment.

#### Hypothesis Testing

Hypothesis 1 aimed to compare interoceptive and exteroceptive attention. To this end we first evaluated a whole-brain interaction between reporting demand [active vs. passive] and attentional target [interoception vs. exteroception] to evaluate whether the effects of attentional target should be evaluated separately for the active and passive reporting conditions. Subsequent analyses compared the simple effects of attentional target within each reporting demand condition, i.e. [Passive Interoception vs. Passive Exteroception] and [Active Exteroception vs. Active Interoception]. To perform these contrasts, all 5 task conditions were estimated separately at the first (individual session) level of analysis and entered into a full factorial design in SPM12.

Hypothesis 2 aimed to investigate whether the differences between exteroception and interoception were moderated by individual differences subjective interoceptive awareness (MAIA scores). Focusing on the contrast of [Active Interoception vs. Active Exteroception], we first created contrast maps at the first (within session) level of analysis. These first level maps were then entered into a second (group) level analysis that included normalized (z-scored) MAIA scale total scores as a covariate of interest. The MAIA covariate was subjected to TFCE correction in the same fashion as other whole brain analyses, and respiration rate change between the two conditions was also included in the factorial model as a nuisance covariate to control for variation associated with changes in respiratory rate.

Hypothesis 3 tested the potential moderating factor of endogenous vs. exogenous control of the respiratory cycle by contrasting each of Active Exteroception and Active Interoception against Active Matching using the same five condition model from testing Hypothesis 1.

Hypothesis 4 aimed to understand how engaging in Active Interoception changes brain connectivity relative to Active Exteroception. To accomplish this aim, a psychophysiological interaction analysis (PPI) was conducted in SPM12 using the Generalized PPI Toolbox (v. 13.1), which improves upon standard PPI analyses by estimating the effect each task condition has upon connectivity independently (McLaren et al., 2012). Here, we used the model employed in Hypotheses 2, a second (group) level full factorial model that contained first level contrasts of [Active Interoception – Active Exteroception] and the MAIA covariate term.

To define a region of interest (ROI), the conjunction of the [Active Exteroception - Active Interoception] contrast and the positive MAIA contrast was evaluated, using a threshold of *p* < .001 for each contrast, resulting in a conjoint probability comparable to conservative FWE correction of *p* < 1 × 10^-6^. The largest cluster observed was used as a seed region, K = 794 voxels, peak Z = 4.06, x = −4, y = 58, z = 14, consistent with dorsal anterior cingulate cortex (Brodmann area 24). At the first level of analysis, mean timecourse activity extracted from this seed region was convolved with separate boxcar regressors indicating the onsets and durations of the Active Exteroception and Active Interoception conditions, and the ensuing whole brain maps were then contrasted to model the PPI effect for each participant session. These first level PPI maps were then collected and analyzed using the same full factorial modelling approach described above.

#### Test-Retest Reliability

The fact that each participant was scanned twice offered a unique opportunity to test the reliability of study effects across two independent scanning sessions. A conjunction analysis for the four main contrasts reported in this paper were conducted. As the TFCE algorithm does not currently perform conjunction analysis, and halving the sample size reduces experimental power, the baseline and post-intervention sessions were each analyzed separately at p < .001^1/2^ = p < .0316, so that the resulting overlap would yield a conjunction p < .001; a minimum cluster size of k ≥ 500 was also applied to each estimate as this cluster size resulted in a FWE corrected p value < .05.

### Data and Code Availability

The full study protocol was pre-registered with the Open Science Framework (https://osf.io/y34ja). All study materials, including behavioral data and fMRI signal extractions, the code to run the experiment and subsequent fMRI analysis, power analysis, statistics, and graphing scripts, are freely available on the Open Science Framework (https://osf.io/ctqrh/).

## Results

### Control Analyses / Manipulation Checks

**Figure 2** summarizes the analysis of the control variables: respiratory frequency, volume/time (RVT), and tracking error. The peak and trough detection algorithm (Figure 2A) allowed for RVT estimation (see Supplementary Materials for similar images of every run in the study).

### Effects of Attention on Respiration Rate and Respiratory Volume / Time (RVT)

The study-wide average respiration frequency was .21 Hz (Table S1). A main effect of task condition was observed, F(4,854) = 9.36, p < .001, with follow-up pairwise comparisons suggesting that respiration rate was slower during Interoception conditions than Exteroception conditions, with an average reduction of .02 Hz, 95% CI [.01, .03] (Figure 2B). However, respiration rate was not slower during than Exteroception conditions in the Active Matching condition (Table S2).

Condition had a large impact on RVT, F(4,854) = 109.7, p < .001 with equivalent respiratory volumes only between the two exteroception conditions (Table S3). Follow-up analyses confirmed that participants breathed more deeply during Passive Interoception than Exteroception, even more deeply during Active Matching, and deepest during Active Interoception (Figure 2C, Table S4).

A frequency / volume tradeoff was also evident, with frequency and RVT negatively correlated within-participant, r(878) = −.452, 95% CI [−.535, −.434], p < .001, which may offset the impact of changing respiration rates on BOLD activity. Nevertheless, to control for the influence of condition on respiration, block-specific respiration frequency and RVT were included as covariates in all second level analyses. Controlling for frequency made had little impact on BOLD activation in subsequent analysis, but greater RVT was associated with widespread cortical deactivation (Figure S3), reinforcing the importance of controlling for RVT during data analysis.

### Interoceptive and Exteroceptive Tracking Accuracy

Participant tracking of the sensory stimuli (respiration and visual circle) was analyzed to in terms of error (in milliseconds) between key presses and the peaks and troughs of stimulus waveforms within each of the active task blocks. Stimulus waveforms were first phase-corrected to maximally match peaks and troughs with keypresses. Error was then measured as the average time difference between stimulus peaks/troughs and key presses (Figure 2D).

Participants accurately and reliably tracked both the respiratory and visual stimuli (Figure 2E, Table S5) with no error differences between breath tracking in Active Interoception, circle tracking in Active Exteroception, or circle tracking in Active Matching (Table S6). Respiration alignment with keypad responses was superior to all other conditions in Active Matching. These results suggest the IEAT successfully matched tracking difficulty between the Active conditions.

### H1: Effects of Attentional and Reporting Demand

A whole-brain interaction analysis between Reporting Demand [Active vs. Passive] and Attentional Target [Exteroception vs. Interoception] implicated the sensorimotor, temporoparietal, and prefrontal cortex (Figure 3A); follow-up analysis of the median signal across all significant voxels revealed deactivation in the Active Interoception and Active Matching conditions relative to the other task conditions (Figure 3B; Table S7).

**Figure 3.**
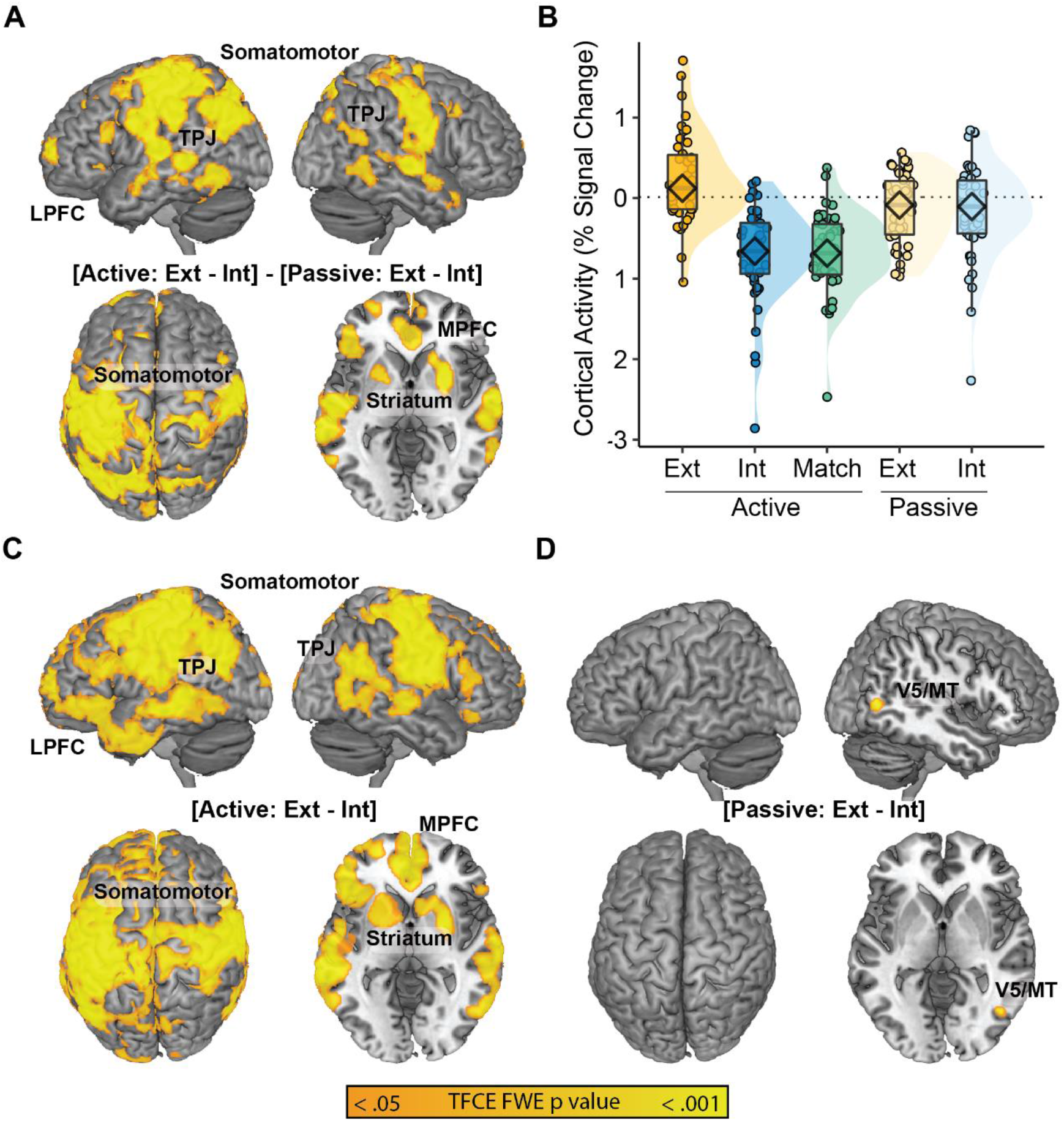
Interactions and Simple Effects of Task **(A)** Interoceptive Reporting Demand [Active vs. Passive] x Attentional Target [Exteroception vs. Interoception] implicated the medial and lateral prefrontal cortices (MPFC and LPFC), somatomotor cortices, striatum, and temporal/occipital cortices. **(B)** Median % signal change across the regions revealed highest activation in the Active Exteroception condition and the two Passive conditions relative to Active Interoception and Act Match. **(C)** Simple effects comparisons between Active Exteroception and Interoception conditions largely replicated the interaction effect, suggesting that the Active conditions were driving the interaction. **(D)** Conversely, only area V5/MT distinguished Passive Exteroception from Passive Interoception.

A follow-up simple effects contrast of [Active Exteroception > Active Interoception] largely replicated the interaction effect (Figure 3C); conversely, the contrast of [Passive Exteroception > Passive Interoception] revealed only a small cluster of activation in motion-related lateral occipital area V5/MT (Figure 3D). The V5/MT finding is consistent with a failure to fully match task features, as Passive Interoception was the only condition in which the circle stimulus remained stationary rather than pulsing. Given a lack of other distinctions between the passive monitoring conditions, subsequent analyses focused upon the active tracking conditions.

### H2: Covariates of Self-Reported Interoceptive Awareness

To better understand the nature of the deactivation observed during Active Interoception relative to Active Exteroception, a focal analysis was conducted to investigate the potential moderating role of self-reported interoceptive awareness (MAIA scores). Interoceptive awareness was significantly related to the level of deactivation observed across a subset of the cortical regions implicated in interoception-related deactivation (Figure 4). Specifically, activity in the anterior cingulate cortex (ACC), dorsomedial prefrontal cortex, and left lateralized language-related regions demonstrated significant associations with interoceptive awareness, such that greater MAIA scores were associated with *reduced deactivation* during Active Interoception relative to Active Exteroception (Table S8). This pattern was also more generally replicated across both passive and Active Matching conditions (Figure S4).

**Figure 4.**
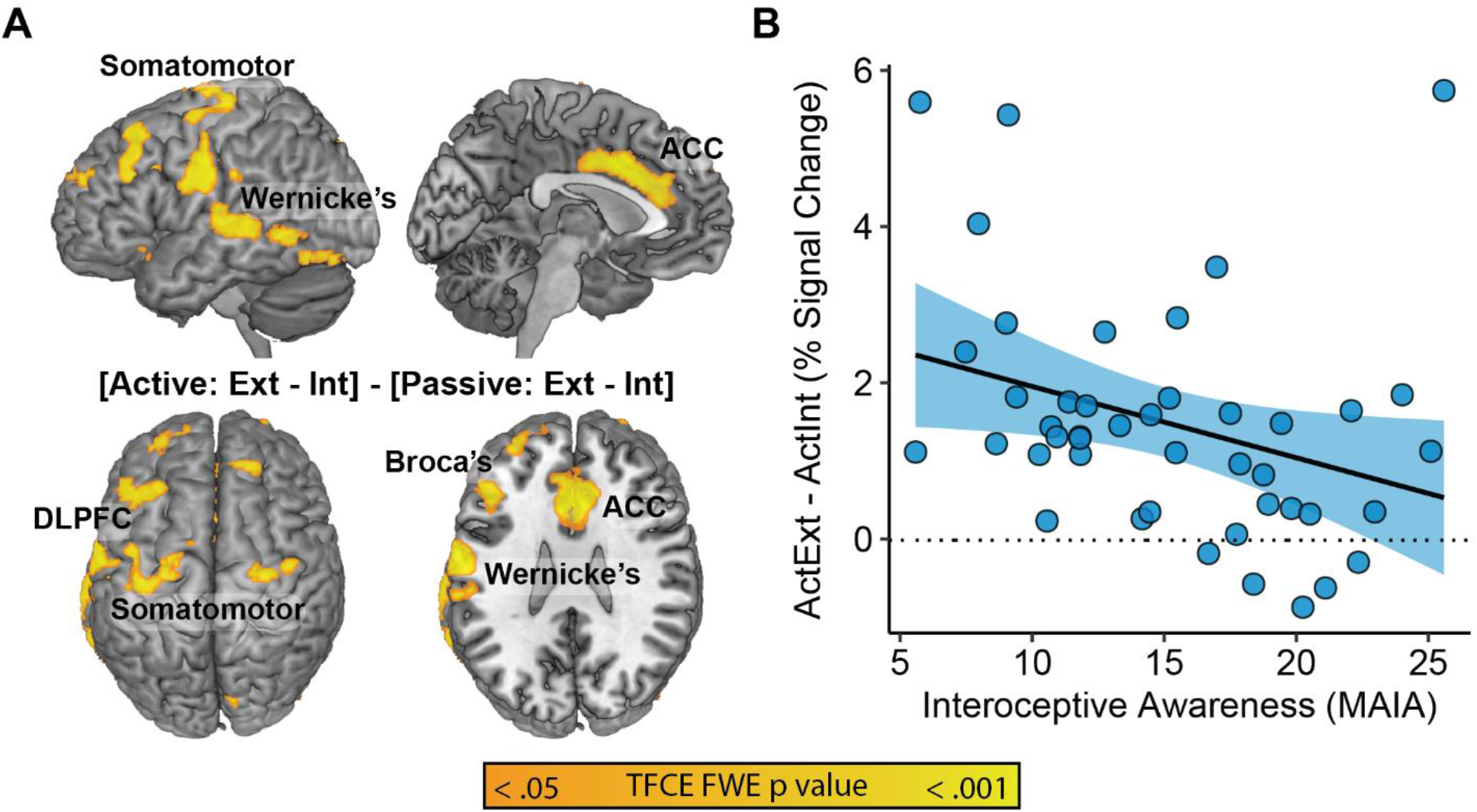
Covariates of Self-Reported Interoceptive Awareness. **(A)** Orange regions denote positive correlates of MAIA scores within the [ActExt – ActInt] contrast. **(B)** Scatterplot of the relationship between MAIA score and median neural activity during the [ActExt – ActInt] contrast across significant covariate regions.

As the largest and most powerfully moderated cluster, the ACC region, k = 1205, x = −4; y = 12; z = 36, was retained as a seed region of interest (ROI) in the PPI analysis (H4) described below.

### H3: Endogenous vs. Exogenous Sources of Interoceptive Control

The next planned comparison contrasted Active Matching against Active Interoception to explore endogenous and exogenous sources of respiratory control. To contextualize Active Matching, we first compared it to Active Exteroception. Active Matching showed even more pronounced and widespread patterns of deactivation than Active Interoception (Figure 3C vs. Figure 5A). Compared to the endogenously paced Active Interoception condition, the exogenously-paced Active Matching condition led to greater deactivation bilaterally along an insula/operculum pathway, somatomotor regions, and within the ventral occipital cortex and cerebellum (Figure 5, Table S9).

**Figure 5.**
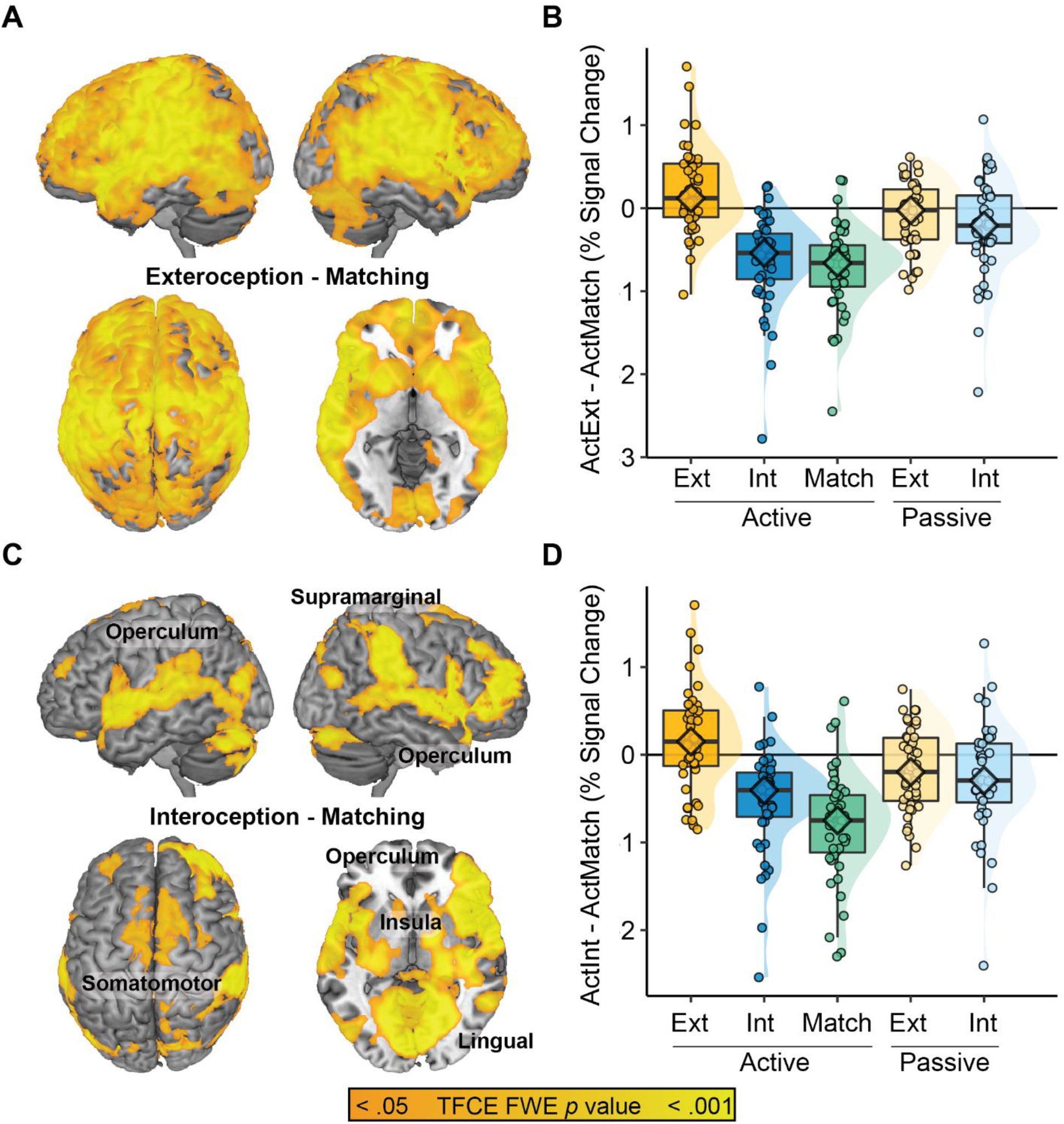
Endogenous vs. Exogenous Sources of Interoceptive Control. **(A)** Active Exteroception > Active Matching replicated the deactivation effect observed for Active Interoception. **(B)** Median % signal change across the regions identified via the [Active Exteroception > Active Matching] contrast. **(C)** Active Interoception > Active Matching linked endogenous respiration to greater activation along an insula-operculum pathway relative to the exogenous control condition. **(D)** Median % signal change across the regions identified via the [Active Interoception > Active Matching] contrast.

### H4: Psychophysiological Interaction (PPI) Analysis

The final contrast examined changes in functional connectivity between Active Exteroception and Active Interoception. The anterior cingulate cortex (ACC) region of interest (Figures 4A & 6A) was entered as a seed region in a psychophysiological interaction (PPI) analysis. The PPI analysis explored changes in functional connectivity with the ACC (Figure 7C) as a function of the two experimental conditions (Active Interoception vs Active Exteroception). The analysis revealed a strong integration of the ACC into regions consistent with the DAN during Active Interoception (Figure 6B; Table S10).

**Figure 6.**
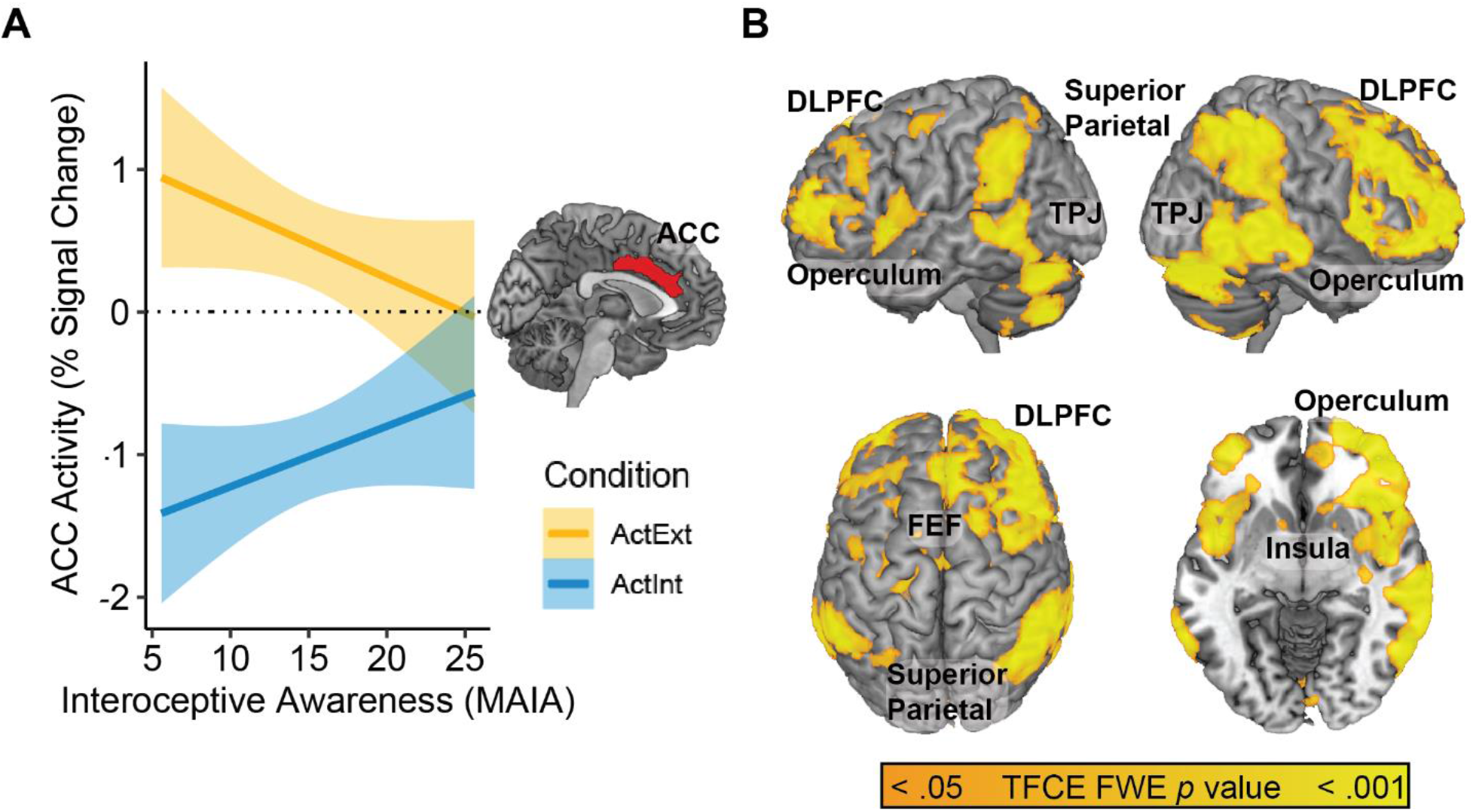
Psychophysiological Interaction (PPI) Analysis. **(A)** The ACC seed region obtained from the MAIA covariate analysis. **(B)** PPI between ACC regional activity and task conditions [Active Interoception > Active Exteroception], demonstrating enhanced ACC connectivity with a frontoparietal network during Active Interoception relative to Active Exteroception.

### Test-Retest Reliability

A conjunction analysis of the main H1-H4 contrasts estimated separately at each of the two timepoints successfully reproduced the results described above (Figure S5).

## Discussion

Relative to visual Exteroception, respiratory Interoception was characterized by pervasive neural deactivation in prefrontal, somatomotor, and temporoparietal regions. This finding deviates from previous respiratory interoception studies (Farb et al., 2013b; Wang et al., 2019), where more difficult sensory tasks yielded greater DAN activity relative to greater DMN activity for less difficult tasks. The finding also differs from liminal heartbeat detection paradigms (c.f., Ring & Brener, 2018), where prior research has related interoceptive accuracy to greater activity and connectivity within the SLN (Chong et al., 2017; Critchley et al., 2004; Tan et al., 2018). The lack of SLN involvement in closely matched, supraliminal tasks supports recent arguments that SLN engagement generally indicates perceptual decision-making rather interoception specifically (Baltazar et al., 2021; Koeppel et al., 2020).

Slower and deeper breathing during the Active Interoception condition could suggest that neural activation differences may be predicated on gross physiological changes driven by different respiration rates, volumes, and spontaneous end-tidal CO2 (PETCO2; Golestani et al., 2015). However, modelling RVT successfully accounted for pervasive deactivation effects (Figure S3), and controlling for respiration rate and RVT at both the first (acquisition volume) and second (task block) levels of analysis failed to eliminate the Interoception deactivation effect. Furthermore, the rate of respiratory slowing (~0.5 Hz) was too small to provoke hypocapnia, and while CO2 was not measured in this study, convergent findings suggest that controlling for PETCO2 reduces functional connectivity but not task amplitude (Golestani & Chen, 2020; Madjar et al., 2012). Finally, the Active Matching task, in which respiration frequency and yoking of behavior were both matched to Active Exteroception, still resulted in this deactivation pattern. While there must certainly be interactions between attention and physiology driving the deactivation effect, they are unlikely to be explained by gross changes in blood oxygenation or CO2.

The right anterior insula, commonly implicated in investigations of interoceptive accuracy (Critchley et al., 2004; Haruki & Ogawa, 2021; Wang et al., 2019), was insensitive to contrasts between actively-reported Interoception and Exteroception. However, the insula was implicated in contrasts between Active Interoception and Active Matching, where moving from endogenous to exogenous respiratory pacing resulted in greater insula deactivation. This effect is not simply due to the integration of external cues, as this deactivation effect was absent in Active Exteroception. Endogenous respiratory rhythms may be continuously processed by this insula/operculum pathway, revealed only when such processing is disrupted by task demands. Here, exogenous control over the respiratory cycle may have disrupted sensory integration or signals such as the body’s internal respiratory ‘clock’, a rhythm commonly attributed to the pre-Bötzinger complex (Smith et al., 1991), but which may communicate with the insula to support homeostasis (Yang & Feldman, 2018). By this logic, heartbeat detection accuracy paradigms implicate the insula precisely because they are comparing participants with robust heartbeat representations to those where such processing is compromised, disrupted, or absent. Conversely, attention to salient signals such as the breath are unlikely to implicate the insula unless compared to situations where breath awareness is somehow disrupted or compromised.

### The Moderating Influence of Interoceptive Sensibility

A second aim of the study was to explore the potential moderating effects of interoceptive sensibility (MAIA scores) on interoceptive network engagement, with the hypothesis that greater subjective interoceptive awareness would be supported by increased SLN activity in the ACC and anterior insula, commensurate with their established role in supporting interoceptive accuracy (Chong et al., 2017; Critchley et al., 2004; Harrison, Köchli, et al., 2021).

Greater activity within the SLN was observed in participants who reported greater interoceptive sensibility. Exactly how the allocation of attention serves to reduce brain activity without impairing performance remains an intriguing question. Performance may be sustained because of noise inhibition along interoceptive pathways (c.f., Kuehn et al., 2016), which could offset the disadvantage seemingly implied by widespread cortical inhibition. This moderating effect is consistent with the ‘low-energy state’ theory proposed above, which characterizes interoceptive attention as a state of preserved attentional monitoring in a less noisy neural environment, especially given findings of increased SLN connectivity during interoceptive attention relative to exteroceptive attention.

### Characterizing the Network Supporting Interoceptive Awareness

A final aim of the study was to characterize an interoceptive attention network supporting adaptive interoceptive sensibility. The rostral ACC was identified as a seed region for psychophysiological interaction (PPI) analysis, convolving the activity in the region implicated as a MAIA covariate with the Active Interoception vs. Active Exteroception contrast.

Interoception led to increased connectivity in the frontoparietal dorsal attention network (DAN; Spreng et al., 2013; Szczepanski et al., 2013). While DAN *activity* was inhibited during interoception, those with greater self-reported interoceptive awareness were likely to offset such deactivation by greater *connectivity* between the ACC and DAN, a potential biomarker of interoceptive engagement.

This finding helps to explain how interoceptive tracking accuracy was equivalent to exteroceptive tracking accuracy despite reduced cortical activity. Interoceptive processing may be continuous and automatic, but largely obscured by a combination of external sensory signals and internal cognitive processing. Interoceptive attention may therefore employ ‘addition by subtraction’, a reduction of competing neural representations rather than the activation of a dormant interoceptive pathway.

The idea of an ‘always on’ interoceptive state is consistent with contemporary theories that place interoception as the background of consciousness, such as Damasio’s “Proto Self” (Bosse et al., 2008) or “Core Consciousness” (Parvizi, 2001), and the distinction between a subcortical “Core Self’ and higher order representations in the cerebral cortex (Northoff & Panksepp, 2008). Interoception is likely the first sense represented in developing brains because of the critical roles homeostasis plays in ensuring survival (Filippetti, 2021; Fotopoulou & Tsakiris, 2017). The consistent presence of periodic interoceptive signals may lead to neural habituation, in keeping with neural repetition suppression effects for familiar and repeatedly represented sensory signals (Barron et al., 2016; Summerfield et al., 2008). Relative to more varied exteroceptive signals, lifelong habituation to omnipresent interoceptive signals may present as widespread cortical deactivation.

### Limitations and Constraints on Generalizability

The IEAT should be improved by introducing a circle motion to the Passive Interoception condition, as the stationary target described here led to a motion confound between passive interoception and the other four experimental conditions. In its current form, Passive Interoception showed relative deactivation in the middle temporal (MT or V5) region of the visual cortex, which is well-established for its sensitivity to motion (Albright & Stoner, 1995).

Several constraints on generalizability are also apparent. We present unexpected and exploratory findings from a relatively small sample; replication is therefore needed, especially given the paucity of studies in this area. The community sample may also not generalize to people with clinical conditions associated with interoceptive dysfunction, nor advanced contemplative practitioners with extensive interoceptive training. Respiration is also only one of many interoceptive signals that each vary in their reportability and controllability, and so may possess distinct neural dynamics. Multimodal interoceptive research is needed to determine the generalizability of the patterns discussed here, to clarify the impact of active reporting vs. passive engagement, the role sensory signal salience, and the influence of participant expertise/dysfunction.

A further limitation to the present findings is that all practices were conducted with eyes-open, which may be dissimilar to many meditation practices. Following repeated evidence that opening one’s eyes decreases the functional connectivity between the SLN and the DMN, it was theorized that reducing the connectivity between these networks reflects an orientation to external rather than internal events (Han et al., 2020). However, here interoceptive attention with eyes open led to greater connectivity between the ACC seed region (an efferent hub of the SLN) and both the DAN and more posterior aspects of the insula rather than the DMN. It remains possible that dynamics for interoceptive awareness may be altered in eyes-closed paradigms. Beyond simply eyes being closed, increased DMN coupling may represent a distinct form of internal awareness at the elaborative, semantic level of processing distinct from interoceptive awareness, as has been previously proposed (Andrews-Hanna et al., 2014; Gusnard et al., 2001).

### Concluding Remarks

Relative to Exteroception, Interoception reduced somatomotor and prefrontal *activity*, offset by enhanced *connectivity* between the SLN and the DAN. Greater interoceptive sensibility was linked to sparing of the SLN and language processing regions from this deactivation pattern.

Together, these processes may allow sensory signals from the body to be better discerned. The ACC deactivation observed in the IEAT paradigm may serve as a candidate biomarker of individual differences in interoceptive processing, one that distinguishes adaptive representations engendered by contemplative training from the avoidance and catastrophizing observed in anxiety and somatic disorders. Larger, more diverse samples and replication of effects are needed to test these emerging ideas.

## Acknowledgements

The content is solely the responsibility of the authors and does not necessarily represent the official views of the National Institutes of Health. We extend our gratitude to Natalie Koh and Sophie Xie for their coordination and data collection efforts, and to Dr. Thomas Grabowski, Dr. Chris Gatenby, and Ms. Liza Young at the Integrated Brain Research Center at the University of Washington for their collaboration. This work was supported by a RIFP award from the School of Nursing at the University of Washington and a grant from the National Center for Advancing Translational Sciences of the National Institutes of Health [UL1 TR002319]. Data analysis infrastructure and trainee salary was supported by a Canadian Natural Sciences and Engineering Council Discovery grant [RGPIN-2015-05901].

