## Supplementary Data for "Interoceptive Awareness of the Breath Preserves Dorsal Attention Network Activity amidst Widespread Cortical Deactivation: A Within-Participant Neuroimaging Study"

**Table S1. Estimated Marginal Means for Respiration Frequency**

| Condition | Mean | SE | df | 95% CI Lower | 95% CI Upper |
|---|---|---|---|---|---|
| Active Interoception | 0.188 | 0.0116 | 24.6 | 0.164 | 0.211 |
| Passive Interoception | 0.197 | 0.0116 | 24.6 | 0.173 | 0.221 |
| Active Matching | 0.209 | 0.0116 | 24.6 | 0.185 | 0.233 |
| Active Exteroception | 0.212 | 0.0116 | 24.6 | 0.188 | 0.235 |
| Passive Exteroception | 0.212 | 0.0116 | 24.6 | 0.189 | 0.236 |

**Table S2. Tukey-Adjusted Pairwise Comparisons of Respiration Frequency**

| Contrast | | | Estimate | SE | df | *t* value | *p* value | |
|---|---|---|---|---|---|---|---|---|
| ActExt | - | ActInt | 0.024 | 0.00507 | 854 | 4.745 | <.0001 | * |
| ActExt | - | ActMatch | 0.002 | 0.00507 | 854 | 0.449 | 0.9916 | |
| ActExt | - | PasExt | -0.001 | 0.00507 | 854 | -0.187 | 0.9997 | |
| ActExt | - | PasInt | 0.015 | 0.00507 | 854 | 2.921 | 0.0294 | * |
| ActInt | - | ActMatch | -0.022 | 0.00507 | 854 | -4.296 | 0.0002 | * |
| ActInt | - | PasExt | -0.025 | 0.00507 | 854 | -4.932 | <.0001 | * |
| ActInt | - | PasInt | -0.009 | 0.00507 | 854 | -1.824 | 0.3601 | |
| ActMatch | - | PasExt | -0.003 | 0.00507 | 854 | -0.636 | 0.9692 | |
| ActMatch | - | PasInt | 0.013 | 0.00507 | 854 | 2.472 | 0.098 | |
| PasExt | - | PasInt | 0.016 | 0.00507 | 854 | 3.108 | 0.0166 | * |

ActExt = Active Exteroception; ActInt = Active Interoception; ActMatch = Active Matching; PasExt = Passive Exteroception; PasInt = Passive Interoception

**Table S3. Estimated Marginal Means for Respiration Volume / Time (RVT)**

| Condition | Mean | SE | df | 95% CI Lower | 95% CI Upper |
|---|---|---|---|---|---|
| Active Exteroception | 2.43 | 0.0407 | 360 | 2.35 | 2.51 |
| Passive Exteroception | 2.51 | 0.0407 | 360 | 2.43 | 2.59 |
| Passive Interoception | 2.72 | 0.0407 | 360 | 2.64 | 2.80 |
| Active Matching | 3.15 | 0.0407 | 360 | 3.07 | 3.23 |
| Active Interoception | 3.42 | 0.0407 | 360 | 3.34 | 3.50 |

**Table S4. Tukey-Adjusted Pairwise Comparisons of Respiration Volume / Time (RVT)**

| Contrast | | | Estimate | SE | df | *t* value | *p* value | |
|---|---|---|---|---|---|---|---|---|
| ActExt | - | ActInt | -0.99 | 0.0573 | 854 | -17.268 | <.0001 | * |
| ActExt | - | ActMatch | -0.713 | 0.0573 | 854 | -12.44 | <.0001 | * |
| ActExt | - | PasExt | -0.078 | 0.0573 | 854 | -1.361 | 0.6527 | |
| ActExt | - | PasInt | -0.289 | 0.0573 | 854 | -5.046 | <.0001 | * |
| ActInt | - | ActMatch | 0.277 | 0.0573 | 854 | 4.828 | <.0001 | * |
| ActInt | - | PasExt | 0.912 | 0.0573 | 854 | 15.907 | <.0001 | * |
| ActInt | - | PasInt | 0.701 | 0.0573 | 854 | 12.222 | <.0001 | * |
| ActMatch | - | PasExt | 0.635 | 0.0573 | 854 | 11.079 | <.0001 | * |
| ActMatch | - | PasInt | 0.424 | 0.0573 | 854 | 7.394 | <.0001 | * |
| PasExt | - | PasInt | -0.211 | 0.0573 | 854 | -3.685 | 0.0023 | * |

ActExt = Active Exteroception; ActInt = Active Interoception; ActMatch = Active Matching; PasExt = Passive Exteroception; PasInt = Passive Interoception

**Table S5. Estimated Marginal Means for Target Tracking Error**

| | | | | | 95% CI | |
|---|---|---|---|---|---|---|
| **Target** | **Condition** | **Mean** | **SE** | **df** | **Lower** | **Upper** |
| Breath | ActMatch | 179 | 28.5 | 28 | 120 | 237 |
| Circle | ActMatch | 231 | 28.5 | 28 | 172 | 289 |
| Circle | ActExt | 239 | 28.5 | 28.1 | 180 | 297 |
| Breath | ActInt | 262 | 28.6 | 28.3 | 204 | 321 |

ActExt = Active Exteroception; ActInt = Active Interoception; ActMatch = Active Matching

**Table S6. Tukey-Adjusted Pairwise Comparisons of Target Tracking Error**

| **Target** | **Condition** | | **Target** | **Condition** | **Estimate** | SE | **df** | ***t* value** | ***p* value** | |
|---|---|---|---|---|---|---|---|---|---|---|
| Circle | Exteroception | *vs.* | Circle | Match | 7.93 | 17.1 | 673 | 0.463 | 0.967 | |
| Breath | Interoception | *vs.* | Circle | Exteroception | 23.68 | 17.3 | 673 | 1.372 | 0.5176 | |
| Breath | Interoception | *vs.* | Circle | Match | 31.61 | 17.2 | 673 | 1.834 | 0.2583 | |
| Breath | Match | *vs.* | Breath | Interoception | -83.79 | 17.2 | 673 | -4.861 | <.0001 | * |
| Breath | Match | *vs.* | Circle | Exteroception | -60.11 | 17.1 | 673 | -3.508 | 0.0027 | * |
| Breath | Match | *vs.* | Circle | Match | -52.17 | 17.1 | 673 | -3.05 | 0.0127 | * |

**Table S7. Task Interactions and Simple Effects**

| Description | Side | Cluster Size (k) | Peak *p* (FWE) | Peak TFCE | Peak Z | Peak *p* (raw) | Co-ordinates (MNI) x | y | z |
|---|---|---|---|---|---|---|---|---|---|
| ***Interaction [ActExt > ActInt] > [PasExt > PasInt]*** | | | | | | | | | |
| Somatomotor, Middle Temporal, Superior Frontal, Temperoparietal Junction | - | 34501 | 0.001 | 2389.64 | 3.54 | 0 | -50 | -32 | 56 |
| Occipital Pole | L | 231 | 0.037 | 1181.47 | 3.24 | 0.001 | -4 | -94 | 30 |
| Angular Gyrus | R | 226 | 0.043 | 1136.62 | 3.16 | 0.001 | 58 | -54 | 20 |
| Insula | L | 169 | 0.043 | 1134.44 | 3.09 | 0.001 | -22 | 8 | -2 |
| Inferior Frontal | R | 70 | 0.046 | 1113.62 | 3.09 | 0.001 | 50 | 24 | 18 |
| ***ActExt > ActInt*** | | | | | | | | | |
| Somatomotor, Middle Temporal, Superior Frontal, Temperoparietal Junction | - | 57088 | 0.001 | 2540.6 | 3.54 | 0 | -50 | -32 | 56 |
| Occipital Pole | L | 318 | 0.032 | 1249.39 | 2.95 | 0.002 | -22 | -94 | 12 |
| Occipital Pole | R | 55 | 0.043 | 1152.73 | 3.16 | 0.001 | 16 | -96 | 16 |
| ***PasExt > PasInt*** | | | | | | | | | |
| Area V5 / MT | R | 62 | 0.033 | 1057.65 | 3.09 | 0.001 | 46 | -64 | 4 |

ActExt = Active Exteroception; ActInt = Active Interoception; ActMatch = Active Matching; PasExt = Passive Exteroception; PasInt = Passive Interoception

**Table S8. Positive Covariates of MAIA Scale on [ActExt > ActInt] Contrast**

| Description | Side | Cluster Size (k) | Peak *p* (FWE) | Peak TFCE | Peak Z | Peak *p* (raw) | Co-ordinates (MNI) x | y | z {mm} |
|---|---|---|---|---|---|---|---|---|---|
| Anterior Cingulate | L | 1205 | 0.025 | 1677.36 | 2.85 | 0.002 | -4 | 12 | 36 |
| Postcentral (Somatosensory) | L | 921 | 0.03 | 1594.48 | 2.79 | 0.003 | -60 | -10 | 32 |
| Precentral (Motor) | R | 122 | 0.036 | 1502.99 | 2.88 | 0.002 | 28 | -14 | 78 |
| Dorsolateral PFC / Broca's Area | L | 489 | 0.037 | 1480.51 | 2.99 | 0.001 | -34 | 32 | 56 |
| Lateral Occipital / Wernicke's Area | L | 1047 | 0.038 | 1467.38 | 2.95 | 0.002 | -52 | -78 | -20 |
| Lateral Occipital | R | 160 | 0.039 | 1464.02 | 2.88 | 0.002 | 54 | -76 | -12 |
| Dorsomedial PFC | R | 61 | 0.042 | 1424.77 | 3.09 | 0.001 | 20 | 46 | 54 |
| Insula | L | 87 | 0.045 | 1386.84 | 2.71 | 0.003 | -42 | -8 | -8 |
| Cerebellum | R | 77 | 0.046 | 1381.72 | 2.91 | 0.002 | 54 | -74 | -36 |
| Cerebellum | R | 74 | 0.047 | 1367.92 | 2.88 | 0.002 | 38 | -44 | -40 |
| Dorsolateral PFC | L | 119 | 0.047 | 1367.11 | 2.79 | 0.003 | -30 | 54 | 30 |
| Lateral Occipital (Superior) | R | 32 | 0.047 | 1362.52 | 2.71 | 0.003 | 10 | -82 | 44 |
| Cerebellum | R | 35 | 0.048 | 1358.1 | 2.95 | 0.002 | 24 | -46 | -50 |
| Frontal Pole | R | 26 | 0.048 | 1355.7 | 2.64 | 0.004 | 26 | 70 | 10 |
| Orbitofrontal | L | 13 | 0.048 | 1351.04 | 2.73 | 0.003 | -56 | 24 | -10 |

**Table S9. Comparisons between Active Effects and Active Match**

| Description | Side | Cluster Size (k) | Peak *p* (FWE) | Peak TFCE | Peak Z | Peak *p* (raw) | Co-ordinates (MNI) x | y | z |
|---|---|---|---|---|---|---|---|---|---|
| ***ActExt > Act Match*** | | | | | | | | | |
| Cerebral Cortex | - | 134079 | 0 | 6116.73 | 3.54 | 0 | -58 | -24 | 16 |
| ***ActInt > Act Match*** | | | | | | | | | |
| Insula, Operculum, Ventral Striatum, Lingual Gyrus | R | 54630 | 0 | 3898.16 | 3.54 | 0 | 60 | 10 | -2 |
| Dorsolateral PFC | L | 706 | 0.011 | 1435.25 | 3.54 | 0 | -32 | 54 | 26 |
| Middle Temporal Gyrus | L | 10 | 0.049 | 952.43 | 3.54 | 0 | -52 | -42 | -6 |

ActExt = Active Exteroception; ActInt = Active Interoception; ActMatch = Active Matching

**Table S10. PPI Effects**

| **<u>PPI: [ActInt - ActExt] * ACC Activity</u>** | | **Cluster** | **Peak *p*** | **Peak** | **Peak** | **Peak *p*** | **Co-ordinates (MNI)** | | |
|---|---|---|---|---|---|---|---|---|---|
| **Description** | **Side** | **Size (k)** | **(FWE)** | **TFCE** | **Z** | **(raw)** | **x** | **Side** | **Size (k)** |
| Dorsal Attention Network: Frontal | R | 23783 | 0 | 3269.41 | 3.54 | 0 | 44 | 24 | 40 |
| Dorsal Attention Network: Parietal | R | 18671 | 0.001 | 2368.12 | 3.54 | 0 | 58 | -40 | 52 |

**Figure S1. Monte Carlo Simulation Results for Study Power Analysis**

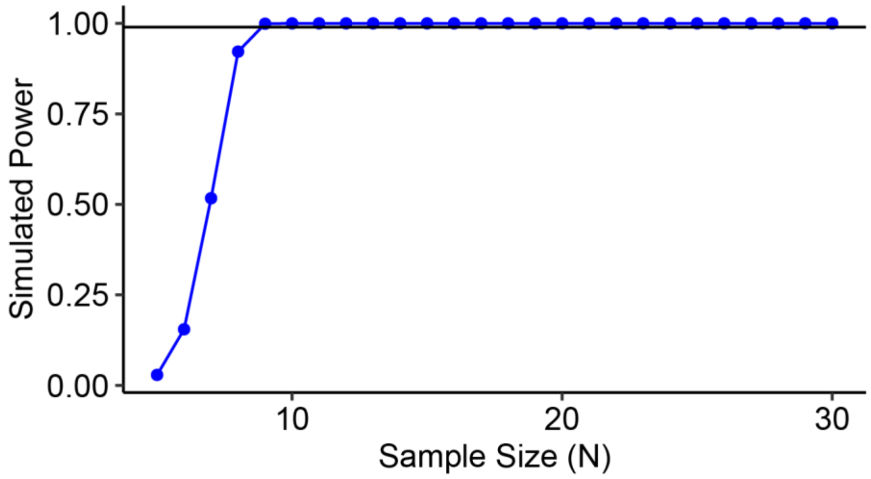

**Figure S2. CONSORT Flow Diagram**

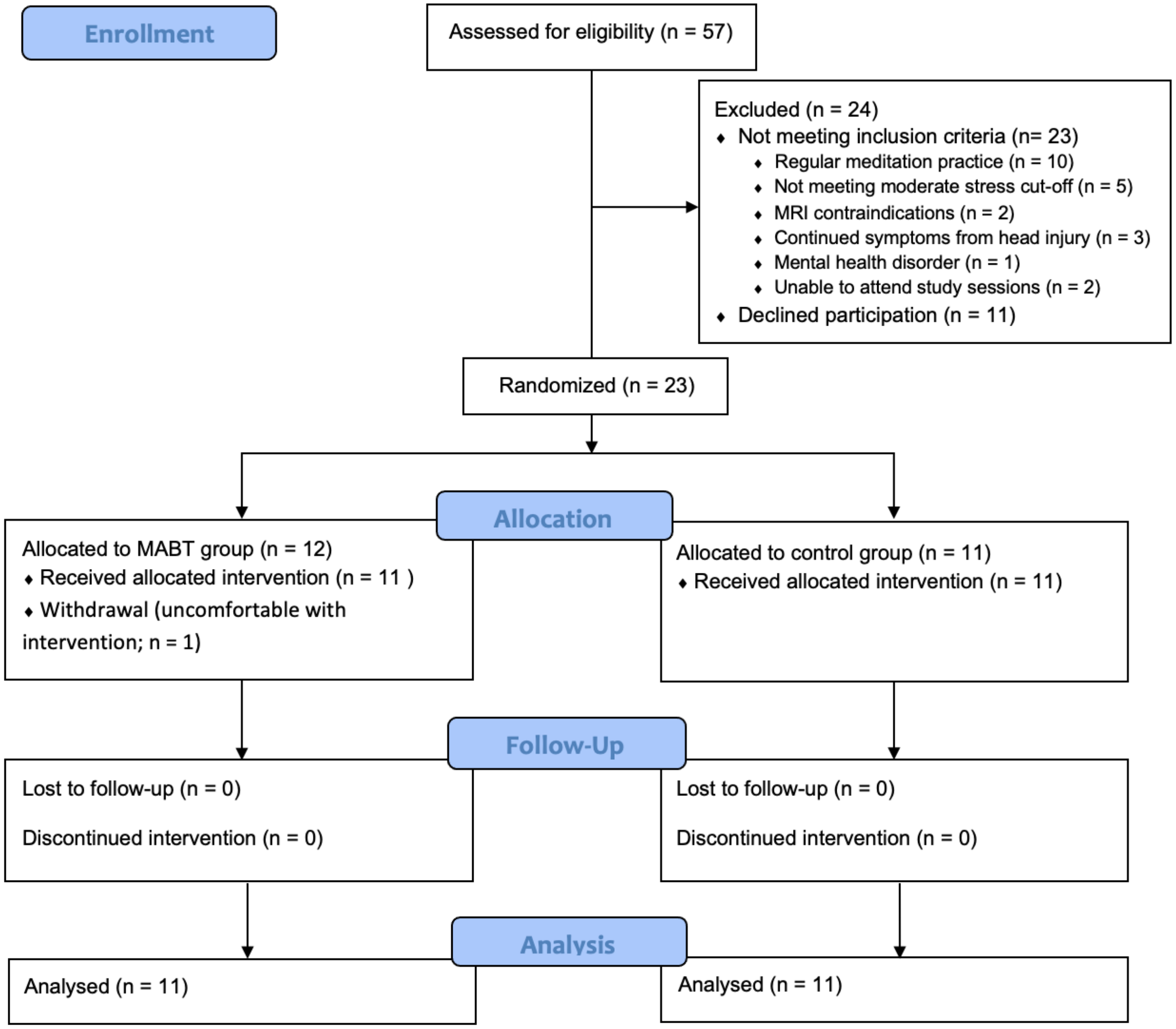

**Figure S3. Negative Contrast of RVT Covariate on BOLD Activity**

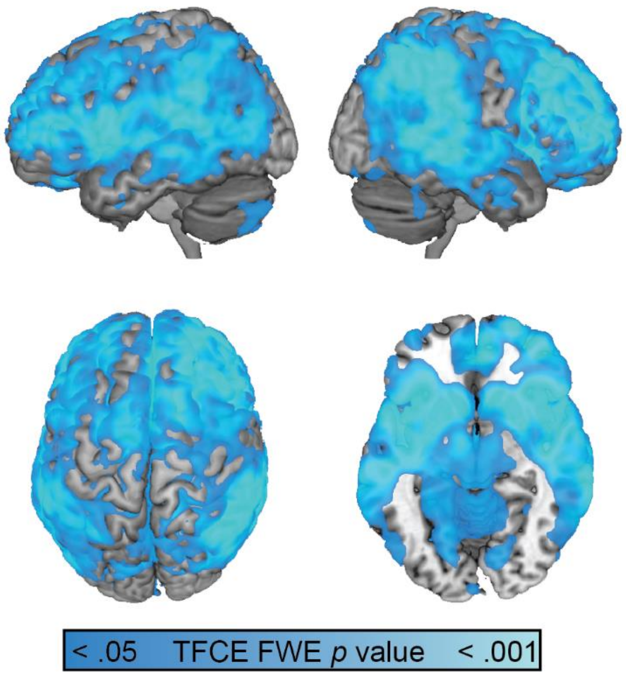

**Figure S4. MAIA Scale and Anterior Cingulate (ACC) Activation with all Five Task Conditions**

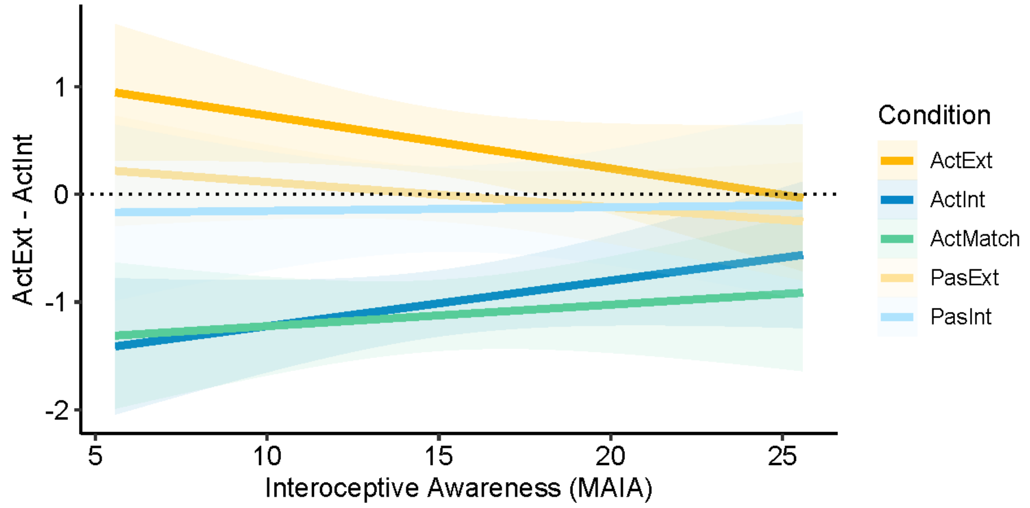


**Figure S4.** The relationship between interoceptive sensibility/awareness scores (MAIA) and Anterior Cingulate activity across all five task conditions. Signal was extracted from each experimental condition and plotted above, demonstrating the same reduced separation between conditions,even within in the passive conditions. ActExt = Active Exteroception, ActInt = Active Interoception, ActMatch = Active Matching, PasExt = Passive Exteroception, PasInt = Passive Interoception.

**Figure 5. Test-Retest Reliability of Neuroimaging Findings**

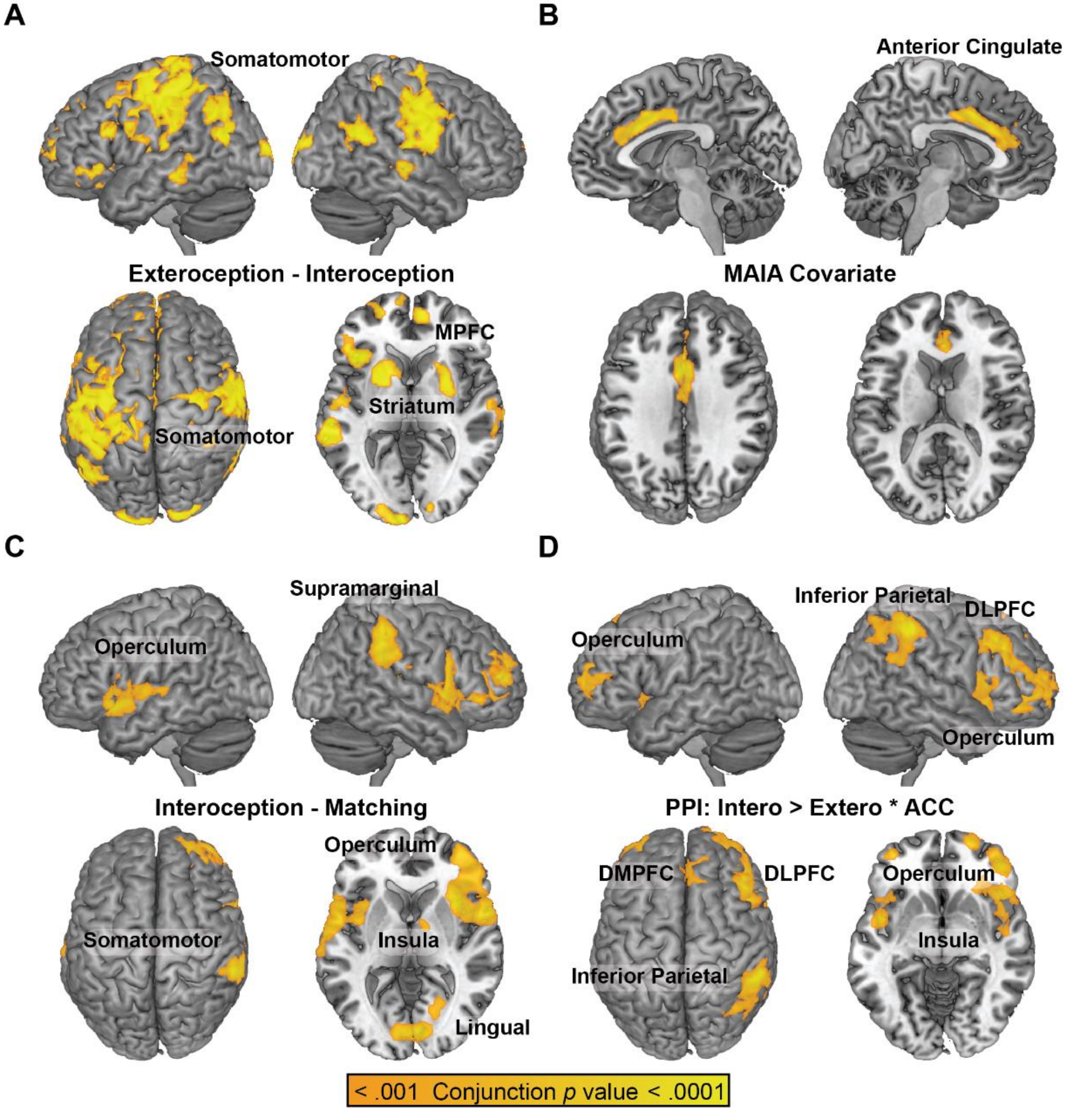


**Figure S5.** To demonstrate the robustness and replicability of the results, conjunction analysis for the four main contrasts reported in this paper were conducted. As the TFCE algorithm does not currently perform conjunction analysis, and halving the sample size reduces experimental power, the baseline and post-intervention sessions were each analyzed separately at $p < .001^{1/2} = p < .0316$, so that the resulting overlap would yield a conjunction $p < .001$; a cluster size of $k \geq 500$ was also applied to each estimate as this cluster size resulted in a FWE corrected p value < .05 for p < .001. **(A)** the contrast of Active Exteroception – Active Interoception reproduced the pattern of somatomotor and prefrontal activation; **(B)** the MAIA covariate of [Active Exteroception – Active Interoception] reproduced the Anterior Cingulate region of interest; **(C)** the contrast of Active Interoception – Active Matching reproduced greater insula/opercular activation for endogenously-paced interoception (Active Interoception) than exogenously-paced interoception (Active Matching). **(D)** The PPI analysis reproduced increased connectivity between the Anterior Cingulate and the dorsal attention network (DAN). DMPFC = dorsomedial prefrontal cortex; DLPFC = dorsolateral prefrontal cortex; ACC = anterior cingulate.
